## Supplemental Information for "Single-molecule analysis of the improved variants of the G-quadruplex recognition protein G4P"

**Protein sequences used in this study:**

The different tags used in the experiment are color-coded, with the FLAG tag represented in orange, 6xHis tag in purple, TEV protease tag in cyan, and thrombin tag in red. Additionally, the RSM elements from RHAU and FANCJ helicases are highlighted in blue and green, respectively.

**G4P** (7.09 KDa)
DYKDDDDKSSGHHHHHHSSGENLYFQGSSG**HPGHLKGREIGMWYAKKQGQKNK**GTGSGAGTGSGAGTGSGA**HPGHLKGREIGMWYAKKQGQKNK**GTGSGA*

**2G4P** (17.32 KDa)
DYKDDDDKSSGHHHHHHSSGENLYFQGSSG**HPGHLKGREIGMWYAKKQGQKNK**GTGSGAGTGSGAGTGSGA**HPGHLKGREIGMWYAKKQGQKNK**GTGSGAGTGSGAGTGSGAGTGSGAGTGSGAGTGSGA**HPGHLKGREIGMWYAKKQGQKNK**GTGSGAGTGSGAGTGSGA**HPGHLKGREIGMWYAKKQGQKNK**GTGSGA*

**FJG4P** (9.93 KDa)
DYKDDDDKSSGHHHHHHSSG**LVPRGS**HM**QDSPEKTTLAAKLSAKKQASIYR**GTGSGAGTGSGAGTGSGA**QDSPEKTTLAAKLSAKKQASIYR**GTGSGA*

**2FJG4P** (18.84 KDa)
DYKDDDDKSSGHHHHHHSSG**LVPRGS**HM**QDSPEKTTLAAKLSAKKQASIYR**GTGSGAGTGSGAGTGSGA**QDSPEKTTLAAKLSAKKQASIYR**GTGSGAGTGSGAGTGSGAGTGSGAGTGSGAGTGSGA**QDSPEKTTLAAKLSAKKQASIYR**GTGSGAGTGSGAGTGSGA**QDSPEKTTLAAKLSAKKQASIYR**GTGSGA*

**Supplementary Table 1. Sequences of DNA used in this study.**

| Name and purpose | Sequence | Structure | Topology |
|---|---|---|---|
| Biotin base strand (for smTIRFM surface tethering) | 5′-GCCTCGCTGCCGTCGCCA-bio-3′ | ssDNA | NA |
| cMycG4 (for smTIRFM surface tethering) | 5′-TGGCGACGGCAGCGAGGCTGGGTGGGTAGGGTGGGTTT -3′ | G4 | Parallel |
| hTelG4 (for smTIRFM surface tethering) | 5′-TGGCGACGGCAGCGAGGCTTAGGGTTAGGGTTAGGGTTTAGGGTTAGGG-3′ | G4 | Mixed |
| 2hTelG4 (for smTIRFM surface tethering) | 5′-TGGCGACGGCAGCGAGGCTTAGGGTTAGGGTTAGGGTTTAGGGTTAGGGTTAGGGTTAGGGTTAGGGTTTAGGGTTAGGG -3′ | G4 | Mixed |
| Cy5-cMycG4 | 5′-/5Cy5/TGGGTGGGTAGGGTGGGTTT-3′ | G4 | Parallel |
| Cy5-hTelG4 | 5′-/5Cy5/TTAGGGTTAGGGTTAGGGTTTAGGGTTAGGG-3′ | G4 | Mixed |
| Cy5-2hTelG4 | 5′-/5Cy5/TTAGGGTTAGGGTTAGGGTTTAGGGTTAGGGTTAGGGTTAGGGTTAGGGTTTAGGGTTAGGG-3′ | G4 | Mixed |
| hTelG4 | 5′-/5Fluor/TTAGGGTTAGGGTTAGGGTTTAGGGTTAGGG -3′ | G4 | Mixed |
| cMycG4 | 5′-/TGGGTGGGTAGGGTGGGTTT3' 6-FAM-3′ | G4 | Parallel |
| Cy5 Biotin base strand (smFRET) | 5′-Cy5-GCCTCGCTGCCGTCGCCA-bio-3 | ssDNA | NA |
| hTel (smFRET) | 5′-TGGCGACGGCAGCGAGGCGGGTTAGGGTTAGGGTTAGGGTTA-Cy3-3′ | G4 | Parallel |

The G4-forming sequences within each oligo are underlined, and sequences complementary to the biotinylated oligo are in blue.

**Supplementary Table 2. Comparison of dissociation constant (Kd) values obtained by different methods and from previous studies.** Note: The numbers obtained from single molecule experiments were adjusted based on the Cy3 labeling efficiency of each respective protein.

| S. No. | Complex (Protein-DNA) | smTIRFM experiments | | | | EMSA Kd (nM) | EMSA Kd Previously published [1] (nM) |
|---|---|---|---|---|---|---|---|
| | | DNA | Protein | Cy3 labeling Efficiency of Protein % | smTIRFM Kd (nM) | | |
| 1. | **G4P-cMycG4** | 100 pM | 150 pM | 55% | 0.22 ± 0.02 | 84.14 ± 8 nM | 2.7 nM |
| 2. | **2G4P-cMycG4** | 100 pM | 100 pM | 71% | 0.16 ± 0.008 | 27.66 ± 4 nM | NA |
| 3. | **G4P-hTelG4** | 100 pM | 200 pM | 55% | 1.6 ± 0.13 | 558 ± 156.8 nM | 14 nM |
| 4. | **2G4P-hTelG4** | 100 pM | 200 pM | 71% | 3.4 ± 0.33 | 86.19 ± 6 nM | NA |
| 5. | **FJG4P-cMycG4** | 100 pM | 200 pM | 42% | 0.27 ± 0.03 | 110.2 ± 58.9 nM | NA |
| 6. | **2FJG4P-cMycG4** | 100 pM | 200 pM | 28.5% | 0.2 ± 0.01 | 130.5 ± 63 nM | NA |
| 7. | **FJG4P-hTelG4** | 100 pM | 5000 pM | 42% | 32.7 ± 4.8 | NA | NA |
| 8. | **2FJG4P-hTelG4** | 100 pM | 2000 pM | 28.5% | 10.7 ± 1.7 | 4526 ± 553 nM | NA |
| 9. | **G4P-2hTelG4** | 100 pM | 200 pM | 55% | 1.9 ± 1.4 | NA | NA |
| 10. | **2G4P-2hTelG4* (3 states)** | 100 pM | 200 pM | 71% | NA | NA | NA |
| 11. | **FJG4P-2hTelG4** | 100 pM | 2000 pM | 42% | 14.5 ± 2.2 | NA | NA |
| 12. | **2FJG4P-2hTelG4** | 100 pM | 1000 pM | 28.5% | 4.6 ± 0.55 | NA | NA |

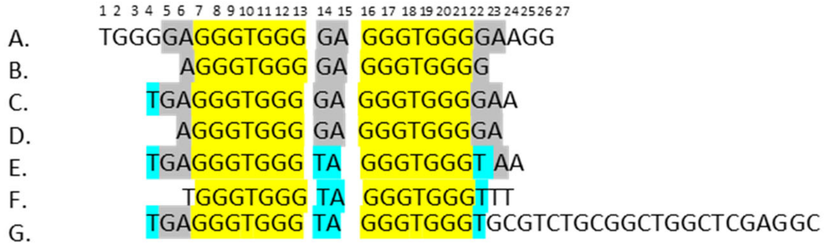


**Supplementary Figure 1**. The promoter sequence of the NHE III1 element of the cMYC gene and its modified versions. A. cMycG4-Pu27: This is the wild-type 27-mer G-rich sequence of the c-Myc NHE III1. B. cMycG4-Pu18: an 18-nucleotide sequence derived from Pu27, which excludes the G2-G5 guanine nucleotides that are not involved in quadruplex formation. C. Myc2345: This is a sequence that can form four major loop-isomers, which can be distinguished by introducing dual G-T mutations at positions 14 and 23, 11 and 23, 14 and 20, and 11 and 20, respectively. These are denoted as 14/23, 11/23, 14/20, and 11/20. D. cMyCG4: This is a shorter version of Myc2345 that was used by Zhen et al. (NAR)[1]. E. cMycG4 14/23: This is a modified version of Myc2345 that forms only one major loop-isomer with the 14/23 mutation. F. cMycG4 14/23*: This is a further modified version of cMycG4 14/23 that was used in this study. G. cMycG4*: This is another modified version of cMycG4 that was used in a previous study published in FASEB[2].

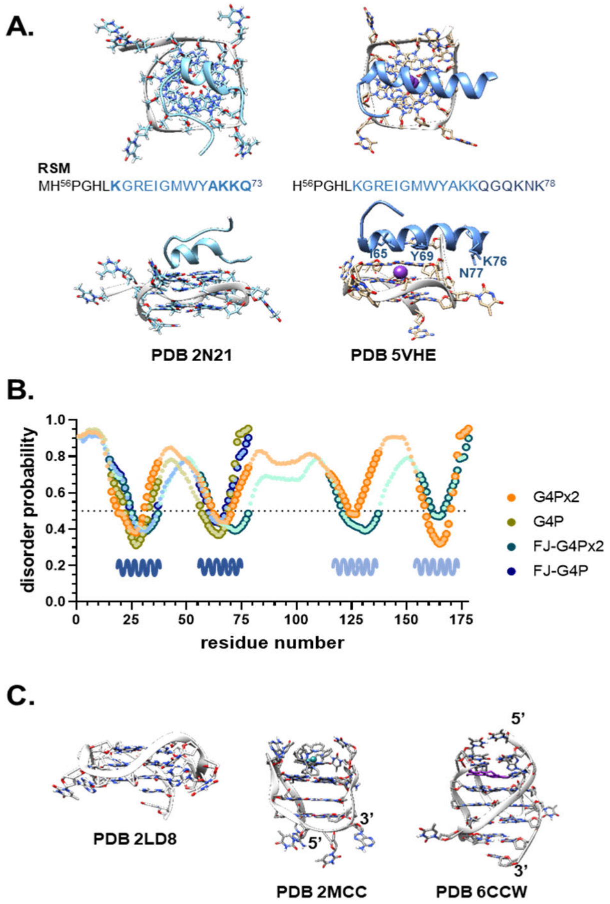


**Supplementary Figure 2. In allG4P constructs, the RSM elements fold into α-helixes connected by disordered regions. A.** NMR structure of the RHAU RSM peptide (PDB 2N21) shows different contacts between the RSM α-helix compared to the longer α-helix containing RSM when a part of RHAU protein (part of the crystal structure from PDB 5VHE). Structures were visualized in UCSF Chimera 1.14. **B**. Disorder prediction for the four constructs using PrDOS server. **C**. Structures of human telomeric G4 in different configurations: 2LD8 – parallel, 2MCC – anti-parallel, 2CCW – hybrid-2.

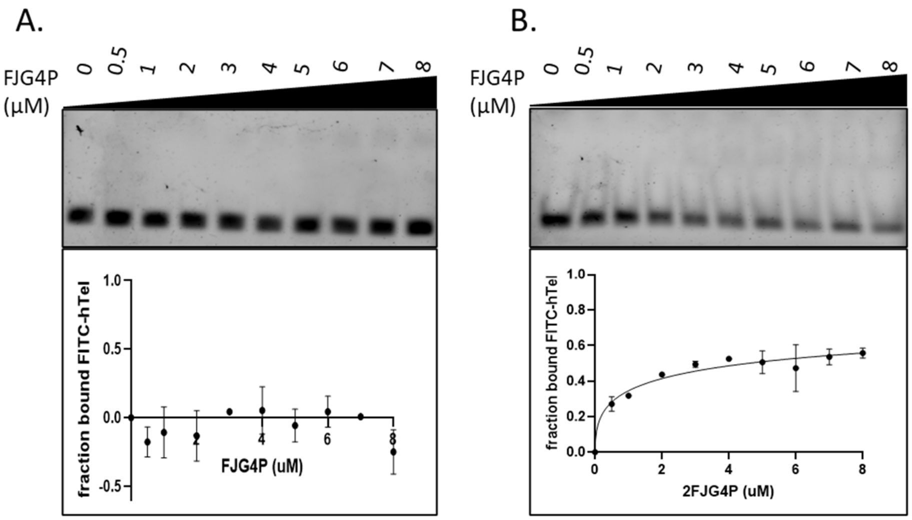


**Supplementary Figure 3. Electrophoretic Mobility Shift Assay (EMSA).** A and B. Indicated concentrations of purified FJG4P and 2FJG4P were incubated with fluorescently-labeled hTelG4 DNA for 10 minutes at room temperature. The samples were resolved on an 8% native polyacrylamide gel and imaged using a ChemiDoc MP imaging system. The bands were quantified using ImageJ software and the data were plotted using GraphPad Prism (n=2).

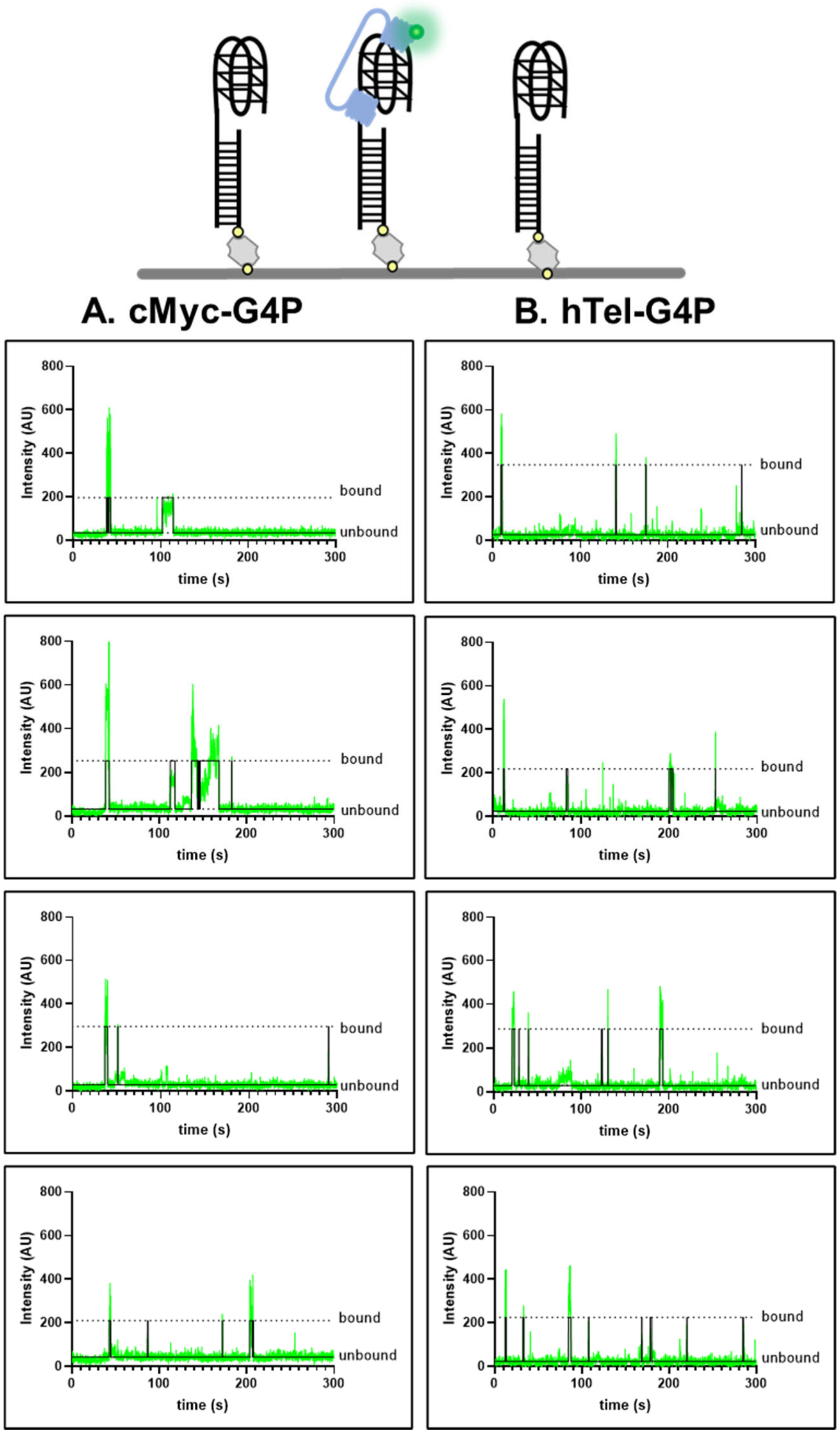


**Supplementary Figure 4. Representative fluorescence trajectories of Cy3-labeled proteins binding to immobilized DNA structures.** Unlabeled DNA was immobilized in the TIRFM reaction chamber, and Cy3-labeled proteins were flown into the chamber. The time-dependent changes in Cy3 fluorescence were recorded, with raw data shown in green and an idealized trajectory shown in black. Panel A and B show the 4 representative fluorescence trajectories for cMycG4-G4P, and hTelG4-G4P each, respectively.

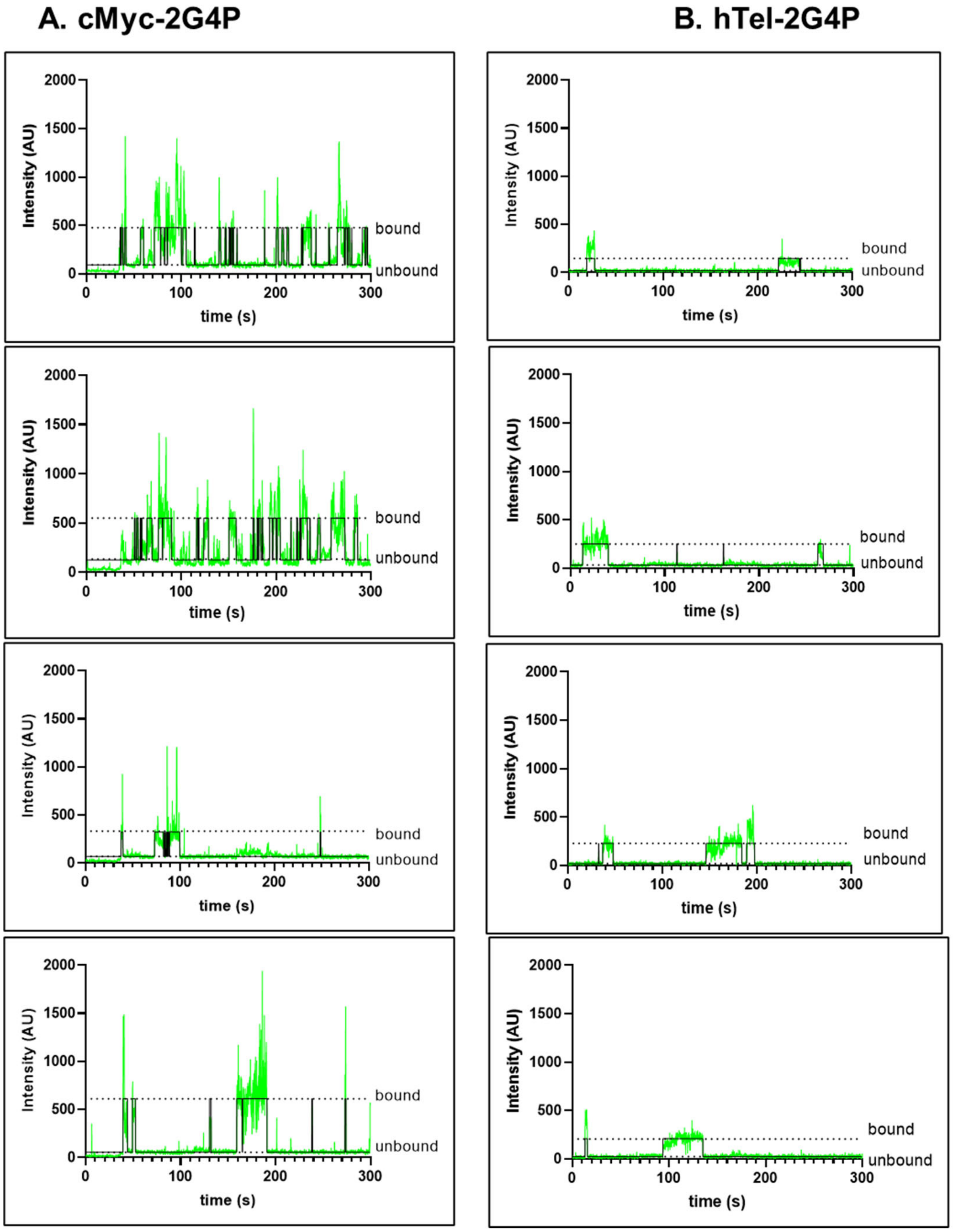


**Supplementary Figure 5. Representative fluorescence trajectories of Cy3-labeled proteins binding to immobilized DNA structures**. Unlabeled DNA was immobilized in the TIRFM reaction chamber, and Cy3-labeled proteins were flown into the chamber. The time-dependent changes in Cy3 fluorescence were recorded, with raw data shown in green and an idealized trajectory shown in black. Panel A and B show the 4 representative fluorescence trajectories for cMycG4-2G4P, and hTelG4-2G4P each, respectively.

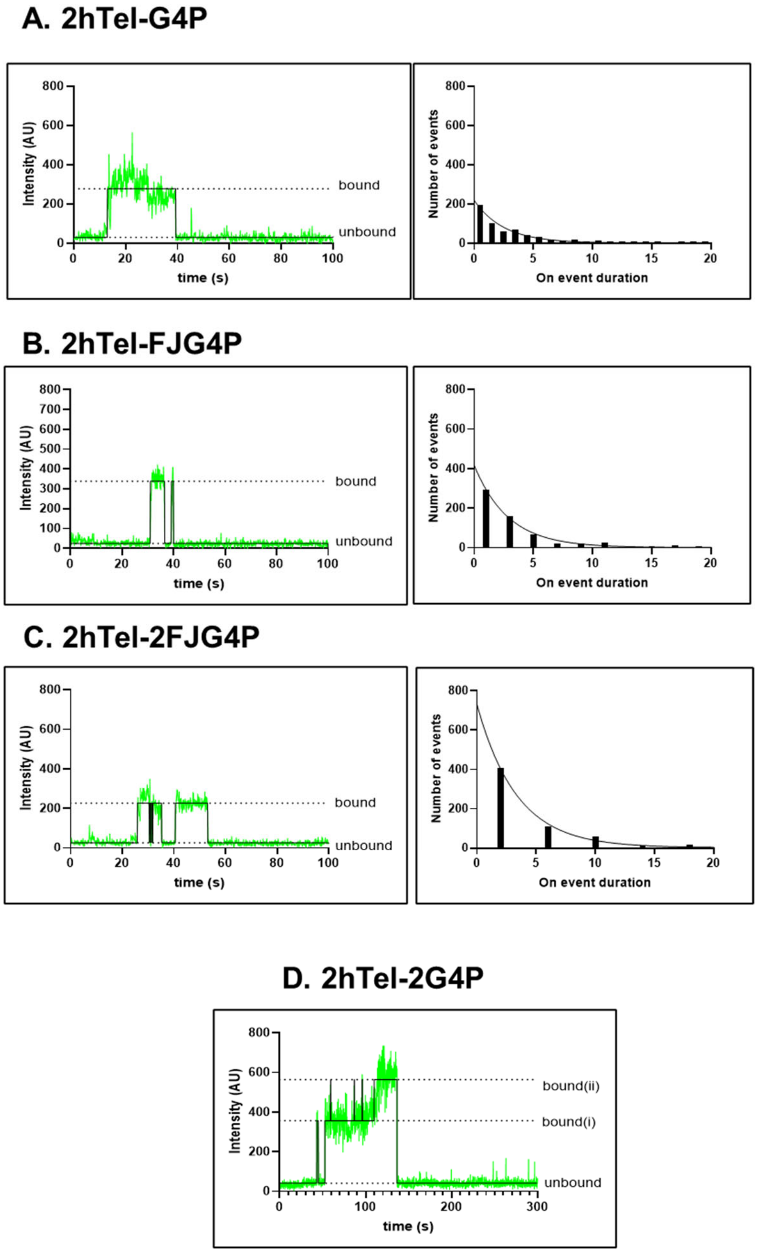


**Supplementary Figure 6. Representative fluorescence trajectories of Cy3-labeled proteins binding to immobilized DNA structures and the dwell-time distribution of all "ON" events**. Unlabeled DNA was immobilized in the TIRFM reaction chamber, and Cy3-labeled proteins were flown into the chamber. The time-dependent changes in Cy3 fluorescence were recorded, with raw data shown in green and an idealized trajectory shown in black. Panel A. 2hTel-G4P, B. 2hTel-FJG4P, C. 2hTel-2FJG4P and D. 2hTel-2G4P showing 3 state binding model.

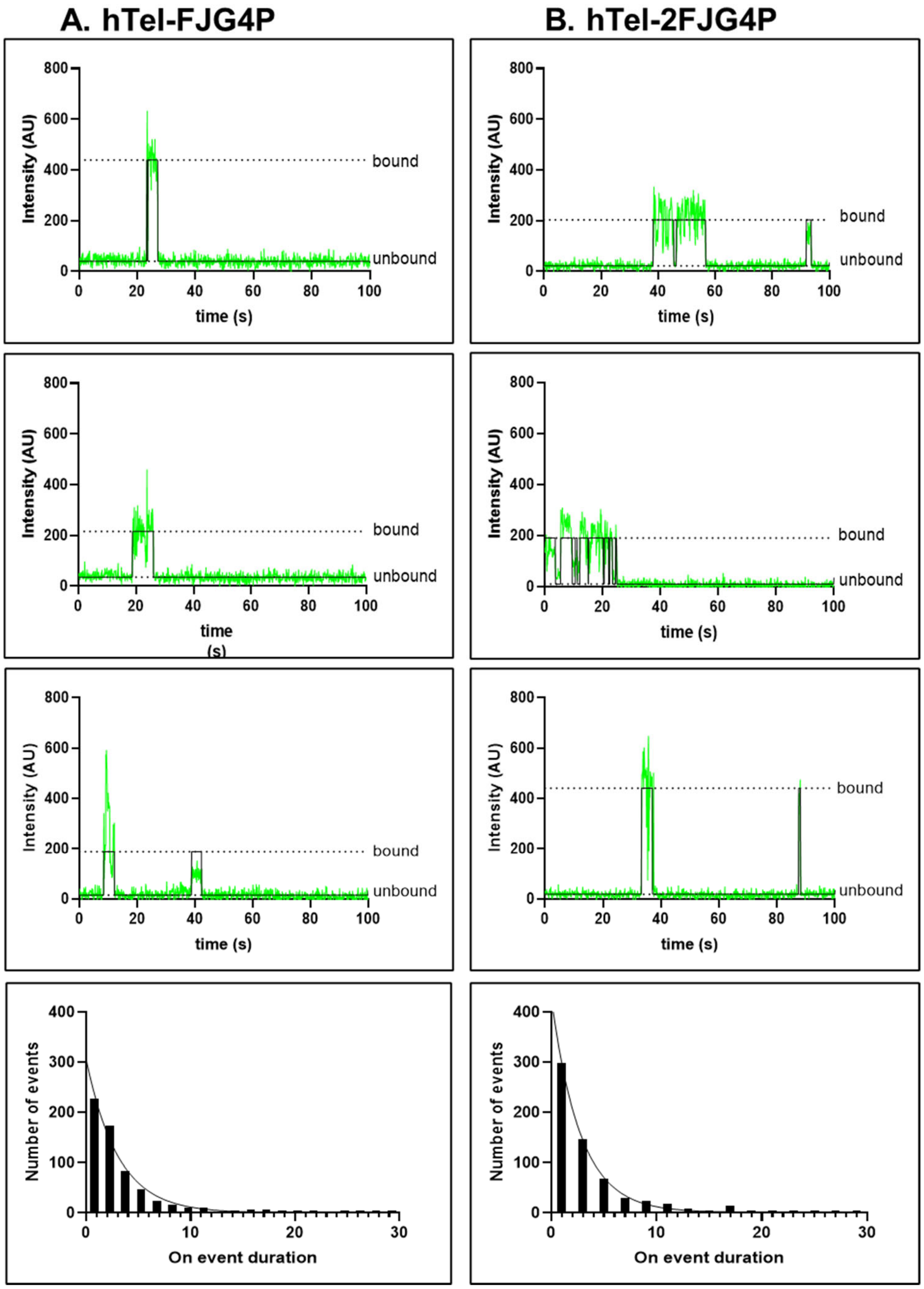


**Supplementary Figure 7. Representative fluorescence trajectories of Cy3-labeled proteins binding to immobilized DNA structures.** Unlabeled DNA was immobilized in the TIRFM reaction chamber, and Cy3-labeled proteins were flown into the chamber. The time-dependent changes in Cy3 fluorescence were recorded, with raw data shown in green and an idealized trajectory shown in black. Panel A and B represent three represented fluorescence trajectories and the dwell-time distribution of all "ON" events for hTelG4-FJG4P and hTelG4-2FJG4P, respectively.

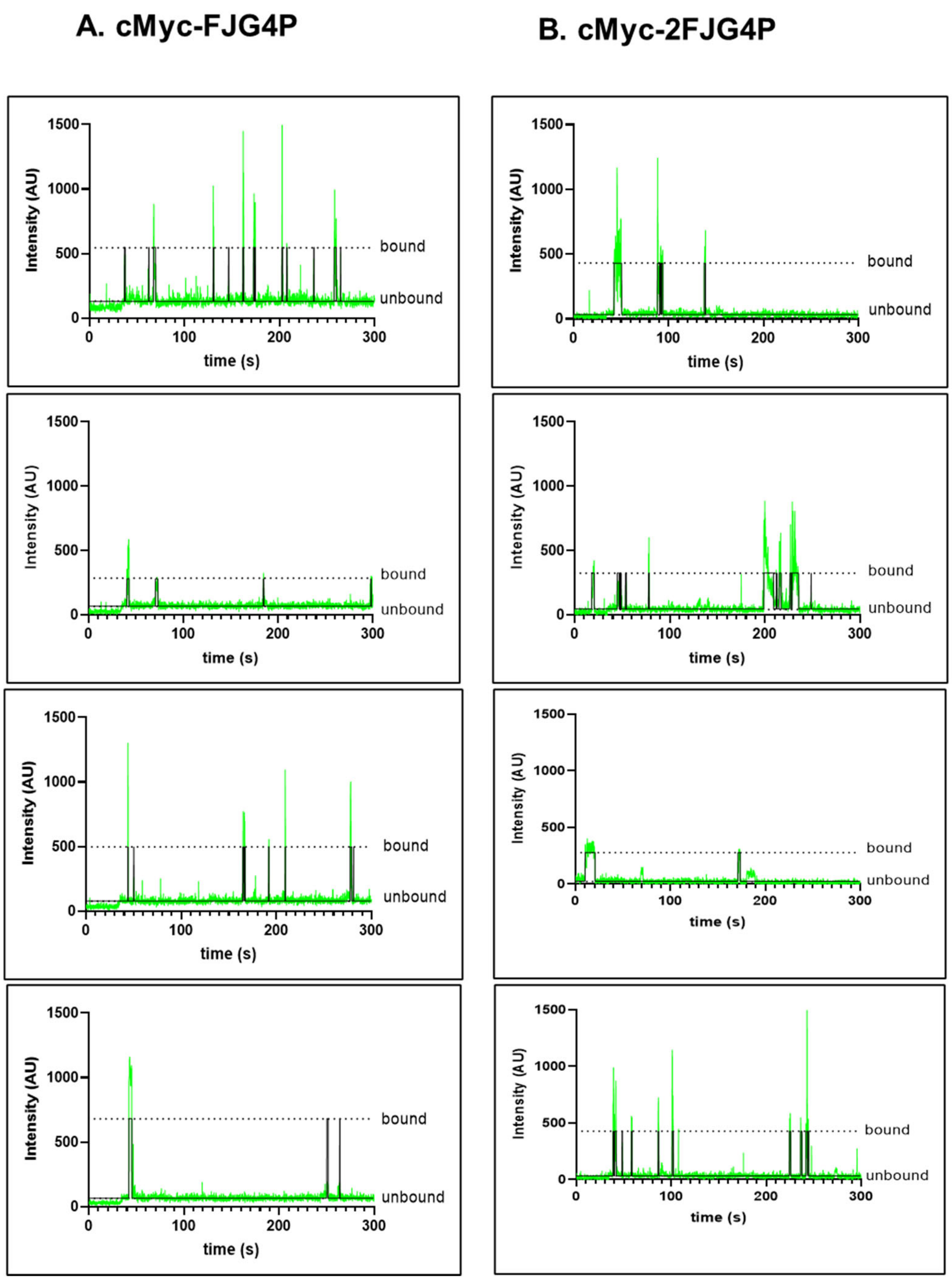


**Supplementary Figure 8. Representative fluorescence trajectories of Cy3-labeled proteins binding to immobilized DNA structures.** Unlabeled DNA was immobilized in the TIRFM reaction chamber, and Cy3-labeled proteins were flown into the chamber. The time-dependent changes in Cy3 fluorescence were recorded, with raw data shown in green and an idealized trajectory shown in black. Panel A and B show the 4 representative fluorescence trajectories for cMycG4-FJG4P, and cMycG4-2FJG4P each, respectively.

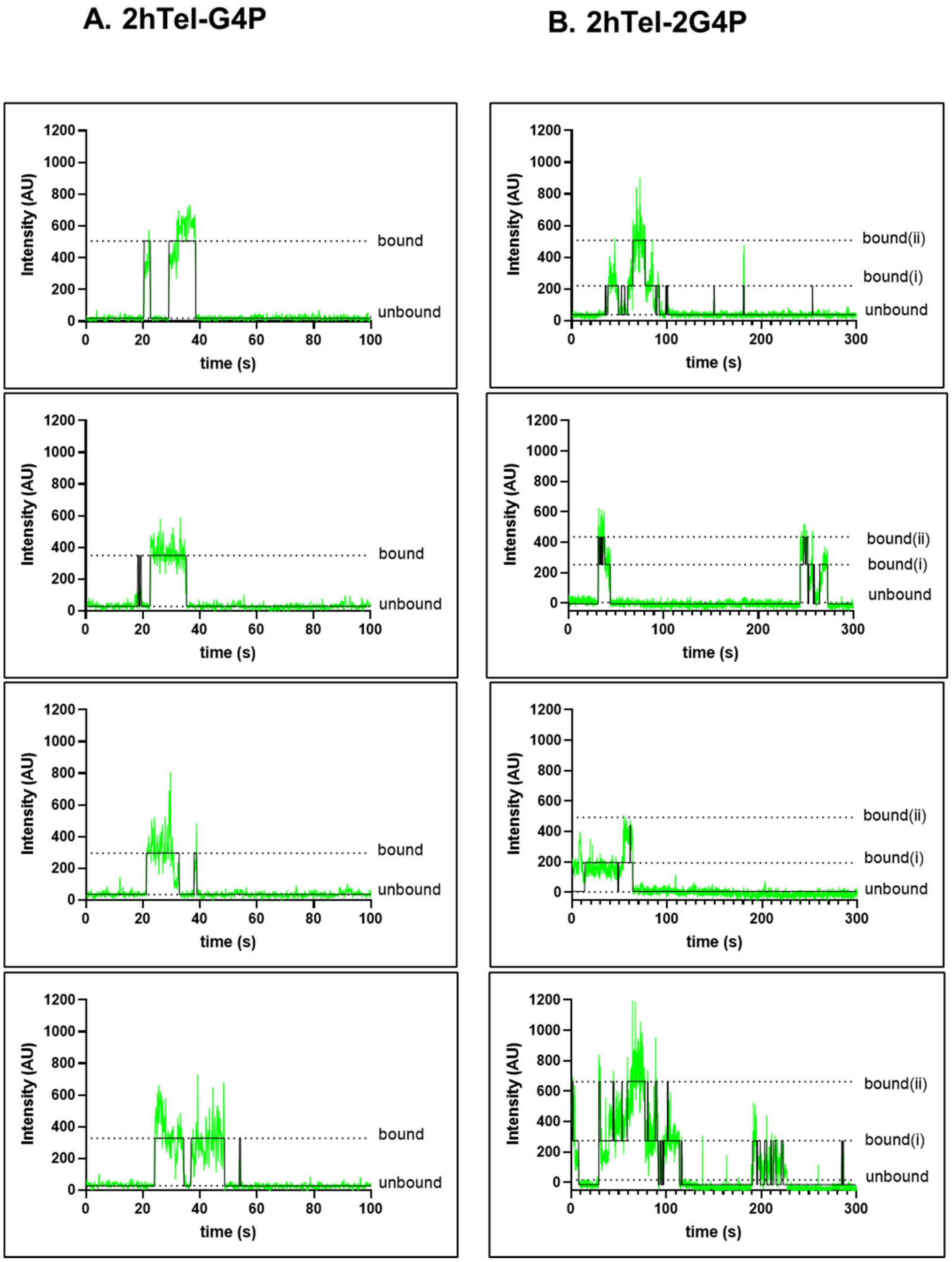


**Supplementary Figure 9. Representative fluorescence trajectories of Cy3-labeled proteins binding to immobilized DNA structures.** Unlabeled DNA was immobilized in the TIRFM reaction chamber, and Cy3-labeled proteins were flown into the chamber. The time-dependent changes in Cy3 fluorescence were recorded, with raw data shown in green and an idealized trajectory shown in black. Panel A and B show the 4 representative fluorescence trajectories for 2hTelG4-G4P, and 2hTelG4-2G4P each, respectively.

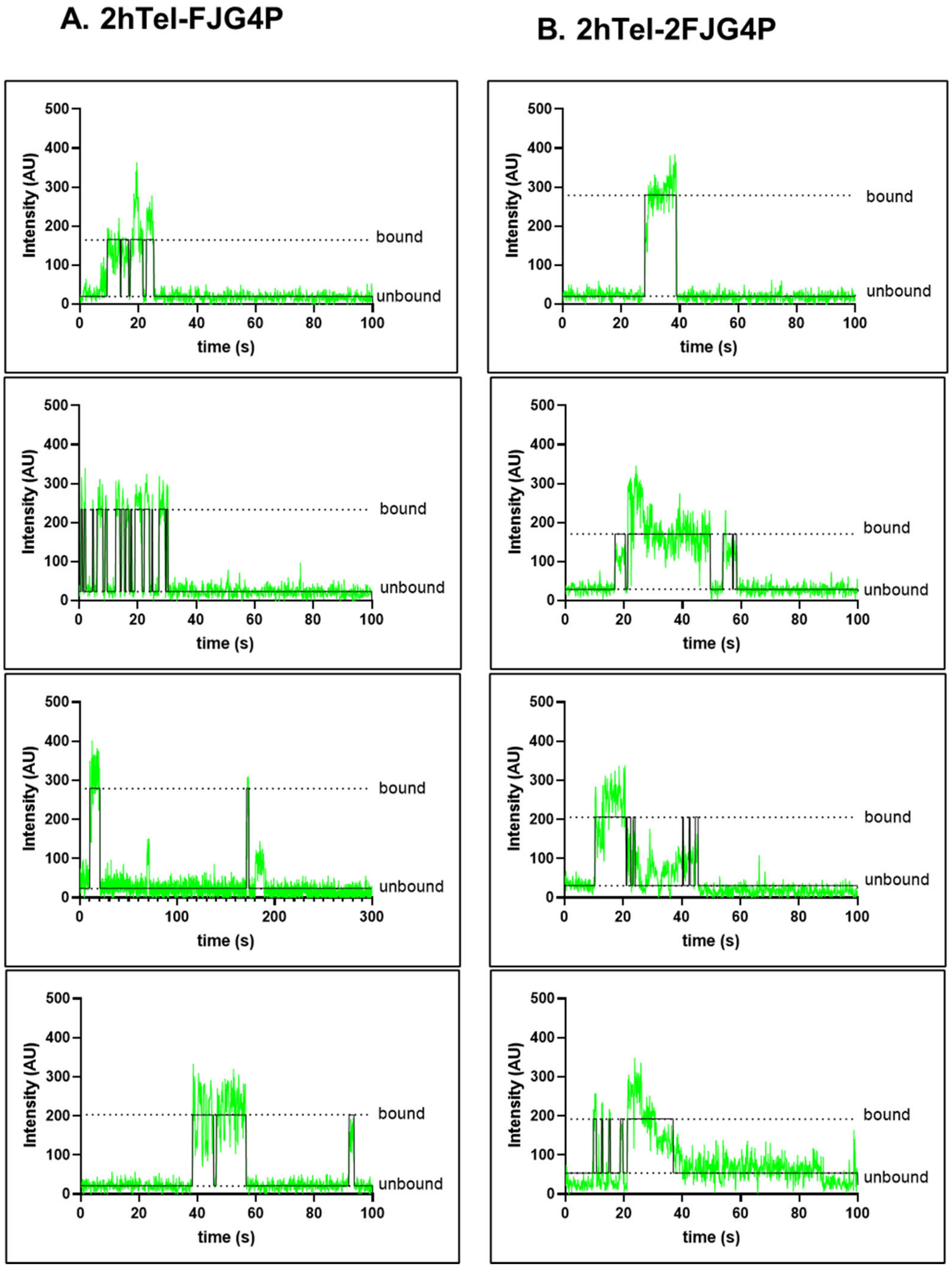


**Supplementary Figure 10. Representative fluorescence trajectories of Cy3-labeled proteins binding to immobilized DNA structures.** Unlabeled DNA was immobilized in the TIRFM reaction chamber, and Cy3-labeled proteins were flown into the chamber. The time-dependent changes in Cy3 fluorescence were recorded, with raw data shown in green and an idealized trajectory shown in black. Panel A and B represent the fluorescence trajectories for 2hTelG4-FJG4P and 2hTelG4-2FJG4P, respectively.

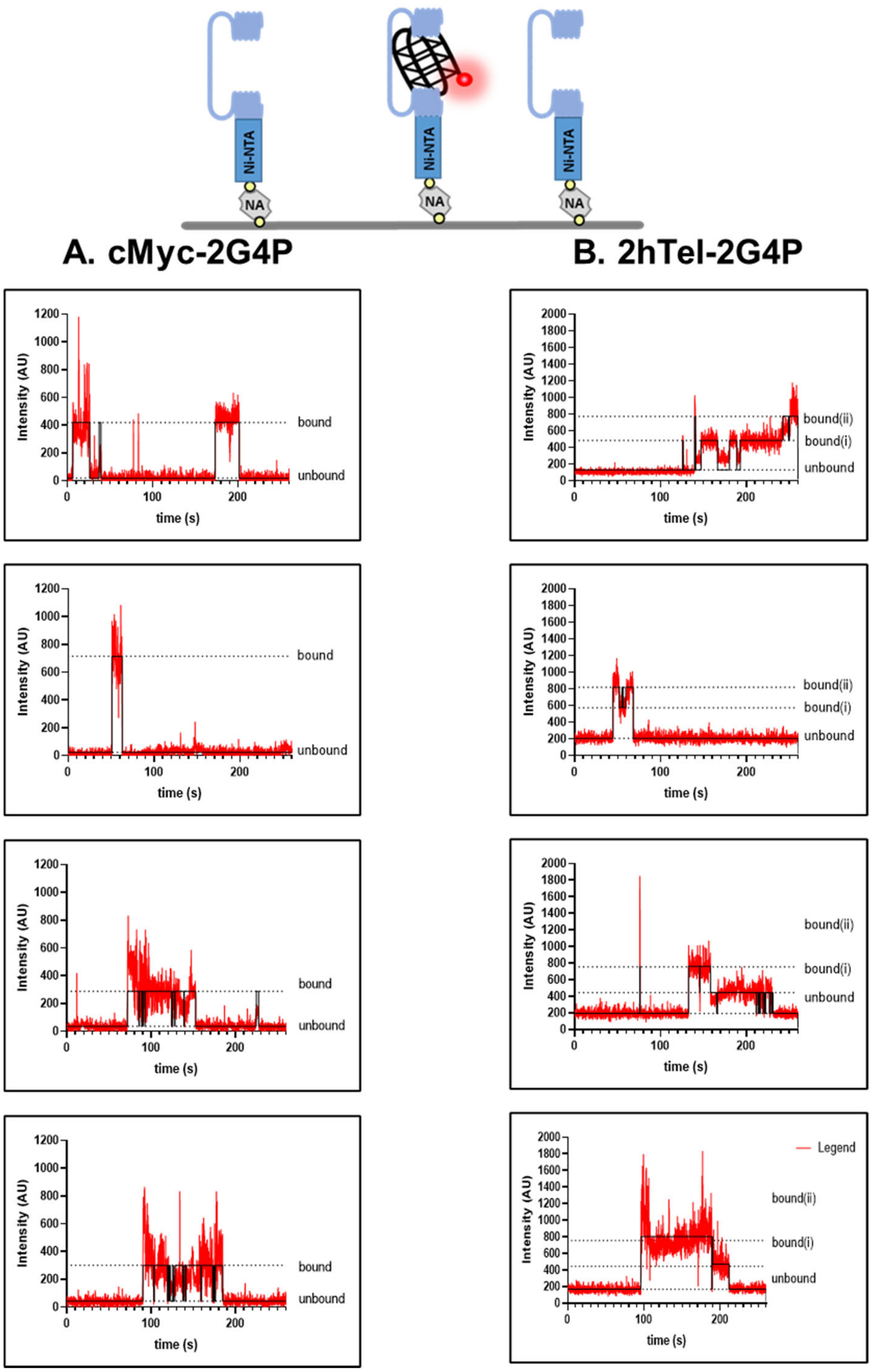


**Supplementary Figure 11. Representative fluorescence trajectories of Cy5-labeled DNA binding to immobilized protein molecules.** Unlabeled protein was immobilized in the TIRFM reaction chamber, and Cy5-labeled DNA molecules were flown into the chamber. The time-dependent changes in Cy5 fluorescence were recorded, with raw data shown in red and an idealized trajectory shown in black. Panel A and B show the 4 representative fluorescence trajectories for cMycG4-2G4P, and 2hTelG4-2G4P each, respectively.

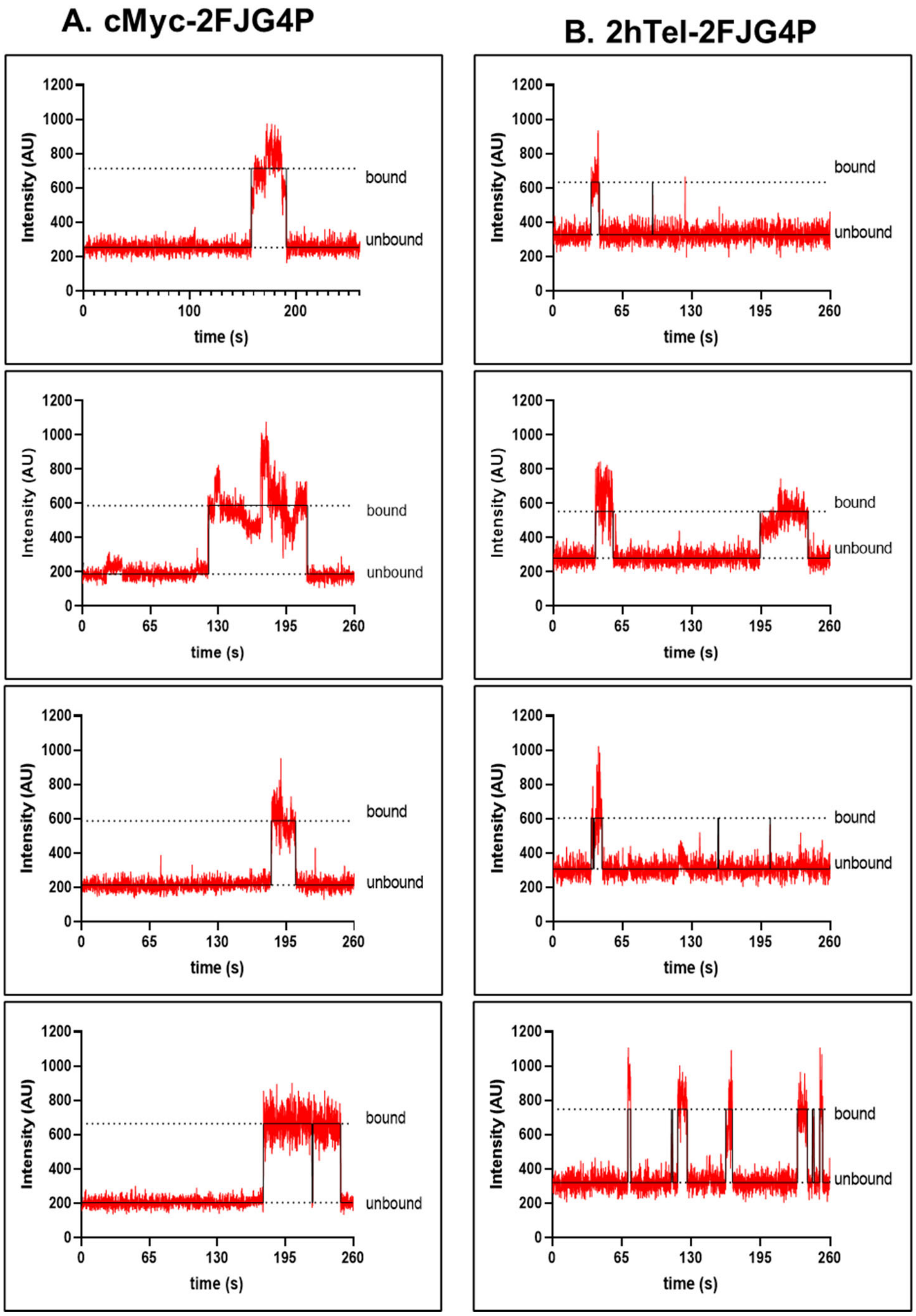


**Supplementary Figure 12. Representative fluorescence trajectories of Cy5-labeled DNA binding to immobilized protein molecules.** Unlabeled protein was immobilized in the TIRFM reaction chamber, and Cy5-labeled DNA molecules were flown into the chamber. The time-dependent changes in Cy5 fluorescence were recorded, with raw data shown in red and an idealized trajectory shown in black. Panel A and B show the 4 representative fluorescence trajectories for cMycG4-2G4P, and 2hTelG4-2G4P each, respectively.

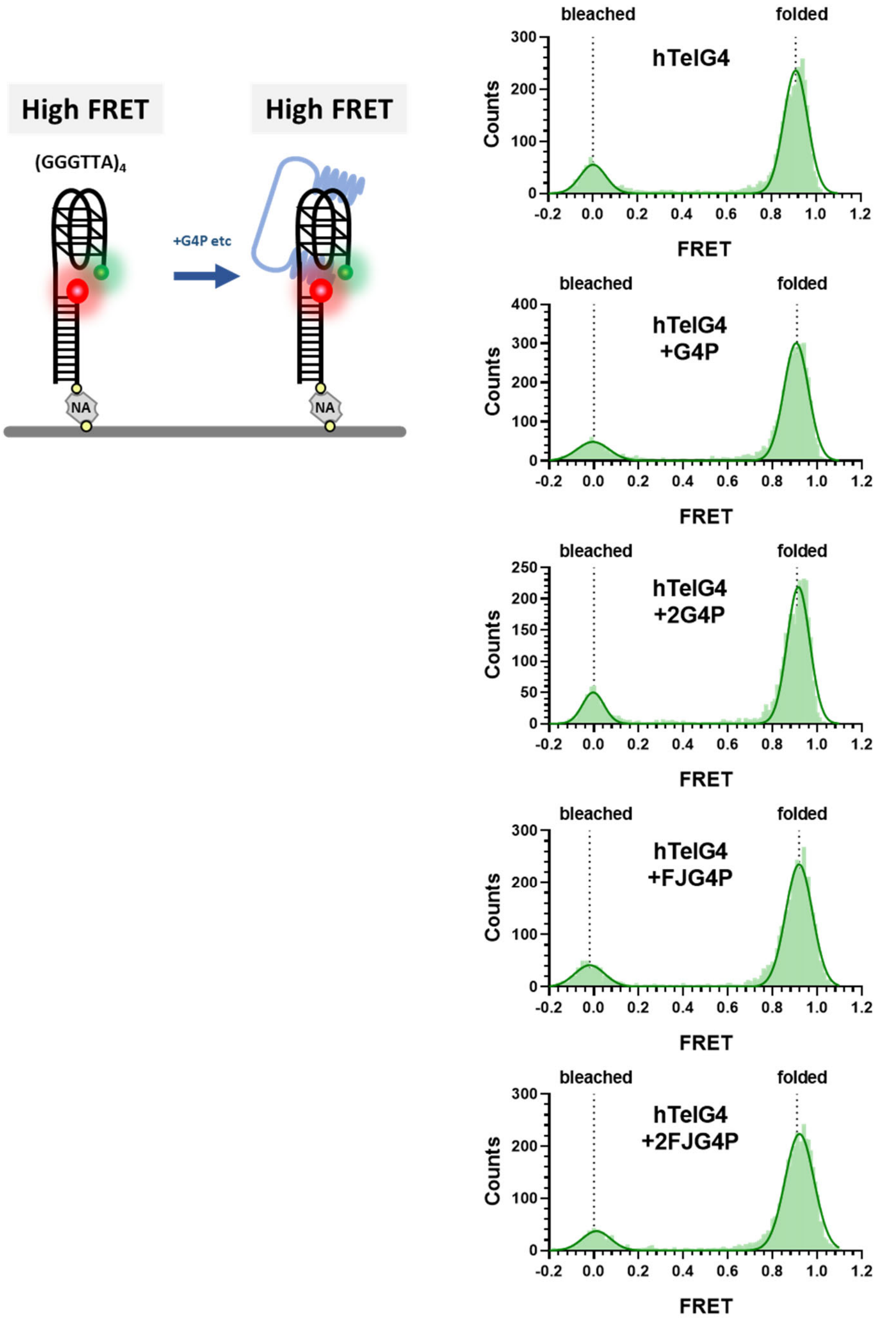


**Supplementary Figure 13. G4P and its variants do not introduce conformational changes to G4 structure.** For all experimental conditions, DNA substrates were immobilized on the surface of the TIRFM flow cell through biotin and Neutravidin (NA)conjugation and incubated with saturation amount of 200 nM G4P, 2G4P and 1 μM of FJG4P, 2FJG4P. Two-color fluorescence trajectories were extracted from short (100 frames) movies. Calculated FRET values for each molecule in the selected movies were binned with the bin size of 0.01 and plotted and fitted to double Gaussian distributions using GraphPad Prism. The resulting FRET distribution histograms are shown. Dotted lines represent means of FRET peaks of the fully folded G4 (folded) and molecules where Cy5 dye was bleached (bleached). The experimental scheme is depicted on the left.

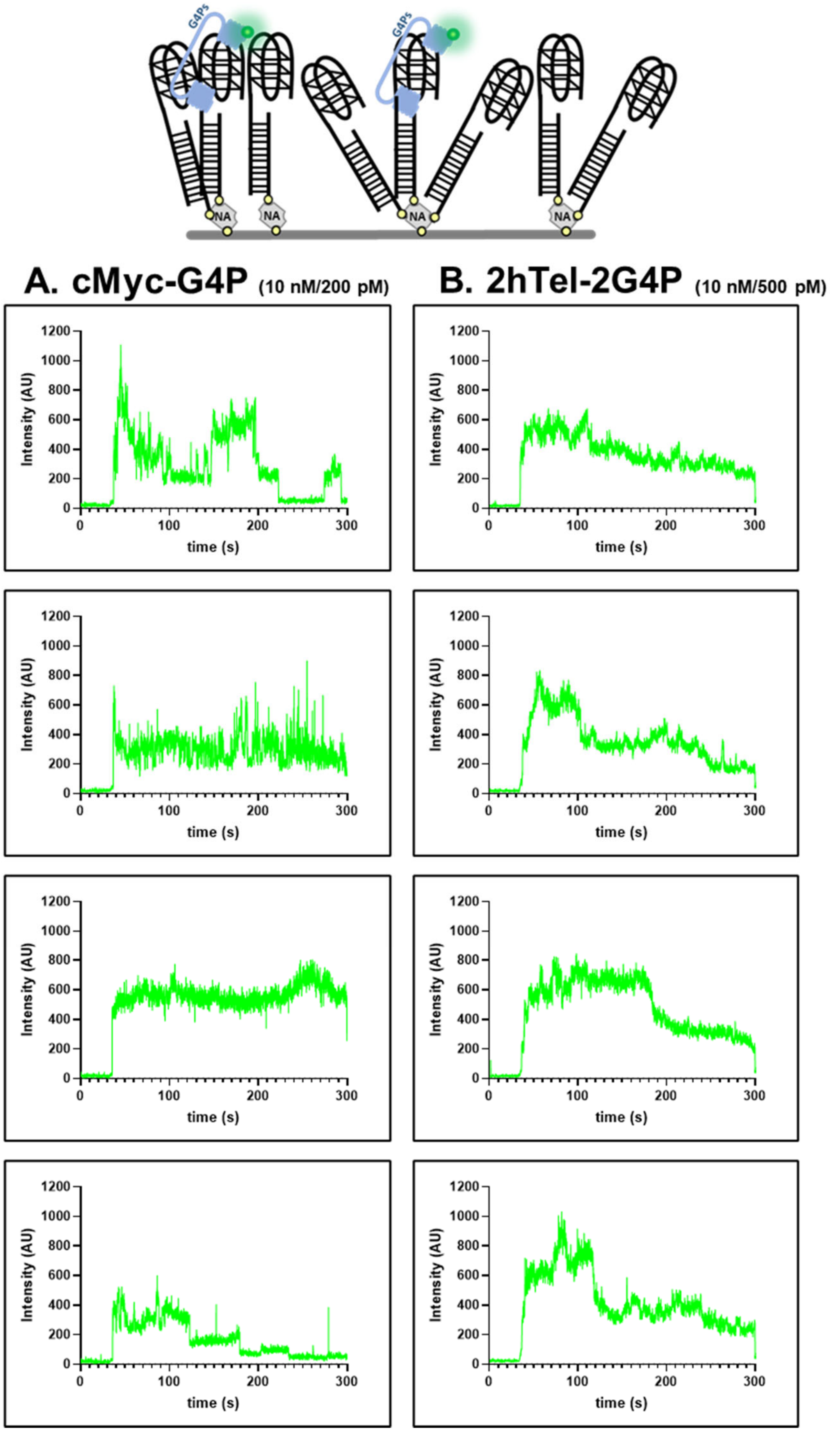


**Supplementary Figure 14. Representative fluorescence trajectories of Cy3-labeled proteins binding to immobilized DNA structures immobilized to the surface at high density** (10 nM). Unlabeled DNA was immobilized in the TIRFM reaction chamber, and Cy3-labeled proteins were flown into the chamber. The time-dependent changes in Cy3 fluorescence were recorded, with raw data shown in green and an idealized trajectory shown in black. Panel A and B represent the fluorescence trajectories for cMycG4-G4P and 2hTel-2G4P, respectively.
